# Illumina-based sequencing framework for accurate detection and mapping of influenza virus defective interfering particle-associated RNAs

**DOI:** 10.1101/440651

**Authors:** Fadi G. Alnaji, Jessica R. Holmes, Gloria Rendon, J. Cristobal Vera, Chris Fields, Brigitte E. Martin, Christopher B. Brooke

## Abstract

The mechanisms and consequences of defective interfering particle (DIP) formation during influenza virus infection remain poorly understood. The development of next generation sequencing (NGS) technologies has made it possible to identify large numbers of DIP-associated sequences, providing a powerful tool to better understand their biological relevance. However, NGS approaches pose numerous technical challenges including the precise identification and mapping of deletion junctions in the presence of frequent mutation and base-calling errors, and the potential for numerous experimental and computational artifacts. Here we detail an Illumina-based sequencing framework and bioinformatics pipeline capable of generating highly accurate and reproducible profiles of DIP-associated junction sequences. We use a combination of simulated and experimental control datasets to optimize pipeline performance and demonstrate the absence of significant artifacts. Finally, we use this optimized pipeline to generate a high-resolution profile of DIP-associated junctions produced during influenza virus infection and demonstrate how this data can provide insight into mechanisms of DIP formation. This work highlights the specific challenges associated with NGS-based detection of DIP-associated sequences, and details the computational and experimental controls required for such studies.

## Importance

Influenza virus defective interfering particles (DIPs) that harbor internal deletions within their genomes occur naturally during infection in humans and cell culture. They have been hypothesized to influence the pathogenicity of the virus; however, their specific function remains elusive. The accurate detection of DIP-associated deletion junctions is crucial for understanding DIP biology but is complicated by an array of technical issue that can bias or confound results. Here we demonstrate a combined experimental and computational framework for detecting DIP-associated deletion junctions using next generation sequencing (NGS). We detail how to validate pipeline performance and provide the bioinformatics pipeline for groups interested in using it. Using this optimized pipeline, we detect hundreds of distinct deletion junctions generated during IAV infection, and use these data to test a long-standing hypothesis concerning the molecular details of DIP formation.

## INTRODUCTION

Influenza A virus (IAV) DIPs were first described over 60 years ago, and are classically defined by their ability to interfere with the production of wild-type virus(1, 2). This ability has been linked to the ability of DI RNAs to both outcompete wild-type (WT) genomic RNAs for resources and packaging into virions, as well as to more potently stimulate the induction of anti-viral immunity through cytosolic RNA sensors (3–6). DIPs have also been implicated in influencing the outcome of influenza virus infection in humans(7). The specific mechanisms and broader functional consequences of DIP formation during IAV infection remain poorly understood.

IAV DIPs are characterized by the presence of large internal deletions in one or more genome segments that disrupt essential open reading frames while retaining the sequences required for replication and packaging(5). As such, the mapping of DIP-associated deletions has helped to define the minimum sequences required for genome replication and packaging (8, 9). These deletions are believed to result from a poorly defined process by which the viral RNA-dependent RNA polymerase (RdRp) ceases RNA polymerization at one site of the viral RNA template (donor site), only to resume at another site downstream (acceptor site), resulting in a failure to copy an internal stretch of the WT template (10). Until recently, the ability to characterize these DIP-associated deletion junction sites (breakpoints) has been limited based on the need to clone and Sanger sequence individual DIP-associated RNAs. As a result, the number of individual DIP-associated RNA sequences that have been analyzed has been relatively small, hindering efforts to define the factors that govern DIP deletion formation.

The advent of next generation sequencing (NGS) has increased the number of individual recombinant sequences that can be identified within a given sample by orders of magnitude. However, the identification and analysis of DIP-associated RNAs by NGS poses new challenges, including the successful alignment of junction-containing (or junction-spanning) reads to the viral reference sequence, the precise definition and localization of DIP-associated deletion breakpoints, and the differentiation of true DIP deletion sequences from the artifactual recombinants that can form during reverse transcription, PCR, and/or sequencing. Without careful optimization and validation, these issues can easily compromise efforts to define the genetic profile of DIP populations.

Here, we describe the development and validation of an Illumina-based sequencing framework for the identification and analysis of influenza virus DIP-associated deletion junctions. The bioinformatics pipeline combines the Bowtie 2 alignment algorithm with the ViReMa (Virus Recombination Mapper) algorithm developed by Andrew Routh and a collection of additional scripts for data processing and analysis (11, 12). We used simulated NGS datasets and a panel of experimental control samples to optimize and quantify the sensitivity, precision, and reproducibility of our pipeline. Subsequently, we used the optimized pipeline to fine-tune the experimental protocol from sample preparation to RNA sequencing to better detect and map DIP-associated deletions generated during experimental IAV infection. This work highlights the computational and experimental controls needed for Illumina-based NGS studies of viral recombination, and provides an optimized, user-friendly sequencing and bioinformatics pipeline for the identification and analysis of IAV DIP-associated sequences. Higher resolution analysis of these deletion sequences can shed light on both the specific molecular mechanisms of DIP formation, as well as how DIPs may affect the overall behavior of viral populations.

## RESULTS

### Overview of the pipeline

The sequencing framework we describe here encompasses sample preparation, sequencing, and data analysis (**Fig 1A**). In brief, we generate 8-segment, full-length amplicons from viral samples and sequence these using the Illumina MiSeq sequencing platform. Datasets are quality-filtered and aligned to the viral reference genome using Bowtie 2 in a conservative manner that disallows soft clipping. Thus, reads containing deletion junctions fail to align, and are fed into the ViReMa algorithm to detect DIP-associated deletion junctions. Finally, the identified junctions are mapped to the viral genome and output as a matrix containing the segment name, junction start and end sites, and NGS read support that can easily be analyzed using additional software tools. Below, we outline the approaches we have taken to optimize and validate the various steps in the process.

**Fig 1.**
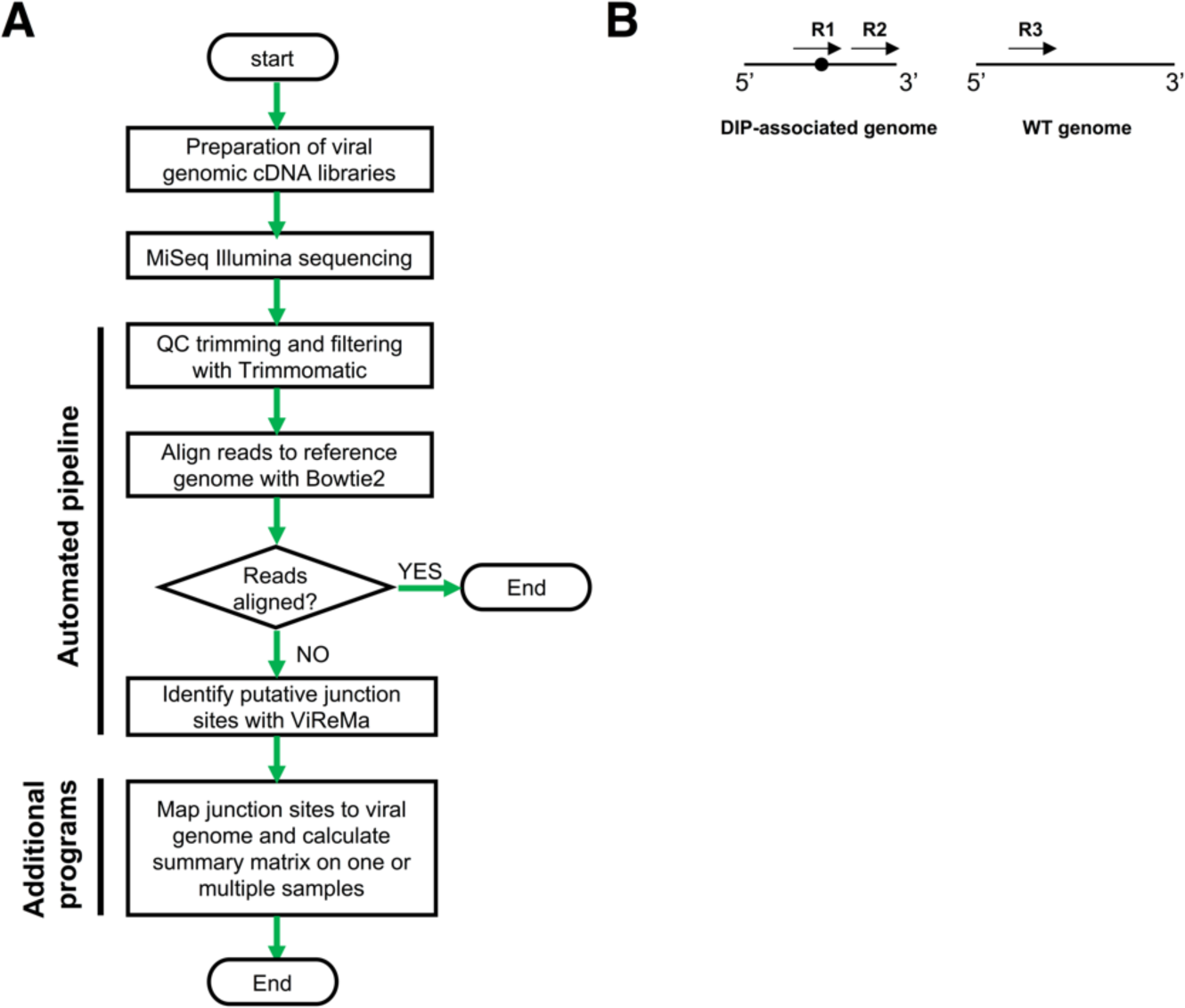
Overview of sequencing/bioinformatics framework. A combination of experimental and computational approaches was extensively optimized in a stepwise manner to establish a pipeline for detecting and analyzing DIP-associated junctions within IAV **(A)** Flowchart outlining the pipeline steps. **(B)** Simple depiction of the possible types of NGS reads in relation to a deletion junction within a sample. Arrows represent individual NGS reads, and the black circle denotes the location of a deletion junction.

### Optimization of analysis pipeline using simulated data

All bioinformatic pipelines have the potential to introduce artifacts and biases during data analysis. Therefore, we first aimed to optimize the sensitivity and precision of our bioinformatics pipeline using simulated NGS datasets where we absolutely know the identity and frequency of all DIP-associated deletion sequences present. IAV DIP-associated deletions can be found in nearly all (if not all) genome segments at a wide range of frequencies (13, 14). To mimic this natural variation, we used MetaSim to generate a panel of Illumina MiSeq-based NGS simulated datasets that contain DIP-associated deletions in all genome segments at varying frequencies and locations (see **Table 1**, **Fig S1**). We used a simple Perl script to randomly generate deletion junctions within the terminal ~600nts of A/California/07/09 (Cal07), since these regions have been shown to be hotspots for DIP-associated deletions(9, 13, 15). We also generated a negative control dataset that lacks deletions to quantify the occurrence of false positives generated by the pipeline. Critically, we introduced a nucleotide substitution frequency of ~1% into these datasets, based on the published Illumina MiSeq empirical error model(16, 17). Each dataset comprised ~1 million 2×250nts paired-end reads, mirroring the read depth that we expect per sample on a typical sequencing run.

**Table 1.** Description of the simulated datasets used in this study.

| Dataset name | Total paired-end reads | Total junction count | Total junction NGS reads | Total WT NGS individual reads | Junction count |  |  |  |  |  |  |  |
| --- | --- | --- | --- | --- | --- | --- | --- | --- | --- | --- | --- | --- |
|  |  |  |  |  | PB2 | PB1 | PA | HA | NP | NA | M | NS |
| Cal07-400 | ~1 million (2X250).<br>Total 2 million individual reads | 400 | 1838898 | 116548 | 50 | 50 | 50 | 50 | 50 | 50 | 50 | 50 |
| Cal07-200 |  | 200 | 1774920 | 225080 | 25 | 25 | 25 | 25 | 25 | 25 | 25 | 25 |
| Cal07 |  | 0 | 0 | 2000000 | 0 | 0 | 0 | 0 | 0 | 0 | 0 | 0 |

### Optimization of alignment

We first optimized the filtering of reads that contain deletion junctions (**Fig 1B**, R1), from those that don’t include junctions (**Fig 1B**, R2,R3). To do this, we aligned all reads to the WT reference genome using Bowtie 2. Reads that successfully align should not contain deletion junctions and are saved for further analysis, while reads that fail to align are fed through the ViReMa algorithm. The performance of this alignment step is highly dependent upon the mismatch penalty scores that are used during alignment. If mismatch penalties are too stringent, reads with random mutations or base calling errors will fail to align and be sent to ViReMa, increasing both the chances of false positives and the total computational time per sample; too lenient, and true junction-spanning reads will successfully align and be excluded from downstream analysis.

We used a junction-rich simulated dataset (Cal07-400) to test the effects of varying the alignment penalty score on the output of ViReMa (**Fig 2A**). We observed that a penalty score of 0.3 minimized the number of unaligned reads (and thus potential for false positives) without diminishing the number junction-spanning reads detected. This value was used for all subsequent analysis.

### Optimization of ViReMa operation

We next optimized the sensitivity and precision with which the pipeline detects deletion junctions. The ability of ViReMa to accurately map true junction-containing reads is affected by three factors. The first is the method the algorithm uses to identify breakpoints. ViReMa extracts and aligns a seed sequence of 20-30nts (the default value of 25 was used in this study) from the beginning of each read and begins aligning the downstream nucleotides. If at any point the downstream alignment fails (as would be the case for a deletion breakpoint), ViReMa generates a new seed sequence starting from that location for realignment. Thus, breakpoints cannot be detected if they occur within the terminal 25 nts of a read.

**Fig 2.**
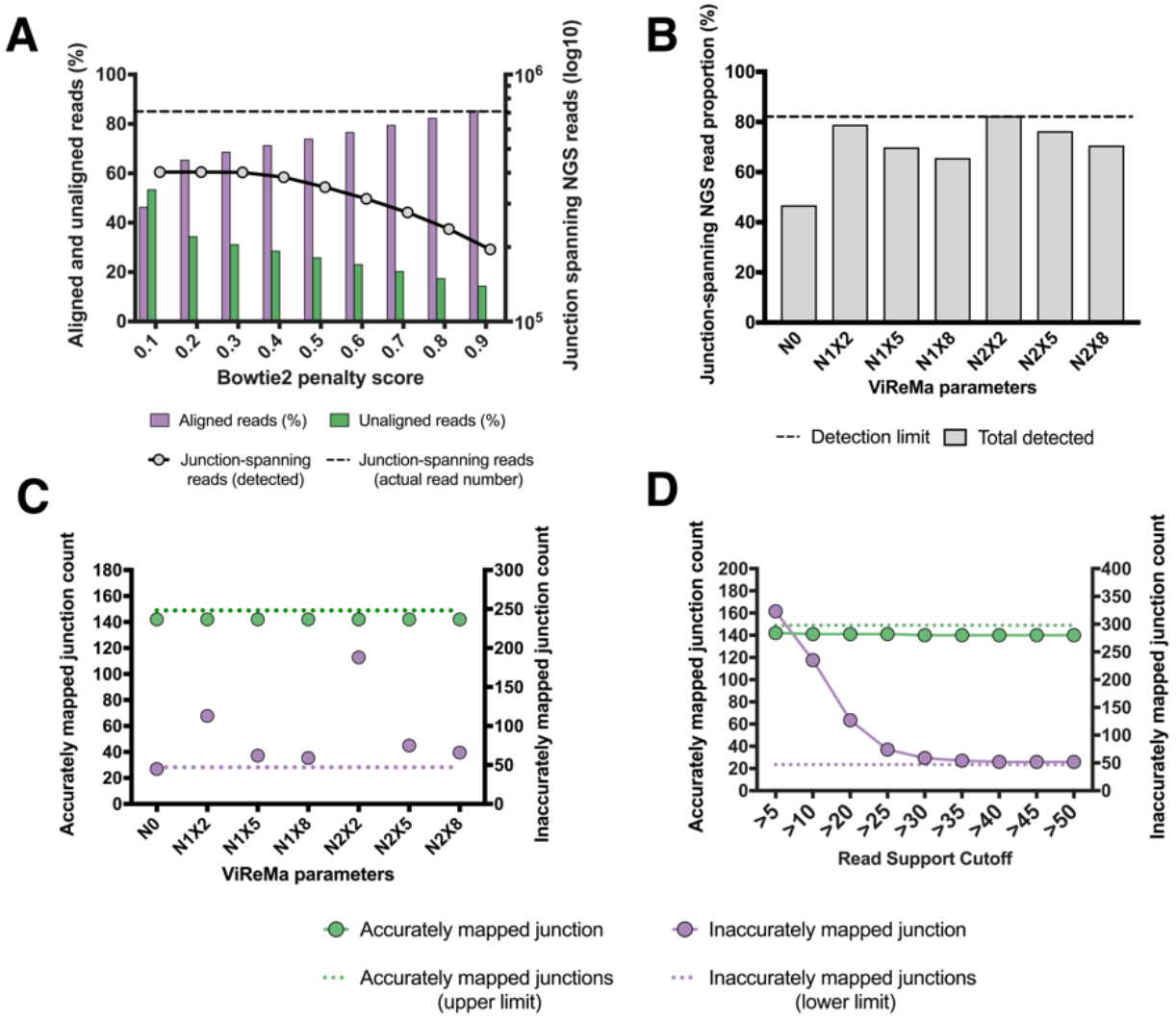
Optimization of bioinformatics pipeline using simulated data. **(A)** Quantification of the effects of varying the bowtie 2 penalty score on the number of junction spanning reads detected by ViReMa in the Cal07-400 simulated dataset (black line; dashed line represents the actual number of junction-spanning reads present in the dataset). The percentages of reads that aligned to reference genome (purple) and failed to align (green) are also shown for each penalty score. **(B)** The effects of ViReMa --X and --N parameters on the percentage of junction spanning reads present in the simulated dataset that were successfully detected. Dashed line shows the maximum theoretical sensitivity (~81.5%), based on the ViReMa seed length of 25nts. **(C)** The effects of ViReMa --X and --N parameters on the number of accurately (green) and inaccurately mapped (purple) deletion junctions reported by the pipeline using the Cal07-200 simulated dataset. The maximum possible number of accurate junctions and the minimum number of inaccurate junctions (resulting from junctions adjacent to direct repeat sequences) are shown for comparison **(D)** Effects of varying the minimum read support cutoff (RSC) on junction detection. Analysis performed on the Cal07-200 simulated dataset using N1X8 ViReMa values.

The second factor is the presence of short direct repeats adjacent to the junction site. These repeats result in a situation where multiple potential breakpoints can give rise to the same final sequence, making precise definition of the true breakpoints impossible (**Fig S2 and S3A**). ViReMa deals with these ‘fuzzy’ regions through the parameter ‘Defuzz’, which can be set to report the junction either to the 5’ end, 3’ end, or the middle of the ambiguous region. For consistency’s sake, we pushed all fuzzy junctions towards the 3’ of the ambiguous region. The effects of direct repeats on breakpoint mapping are impossible to avoid and vary somewhat between IAV genome segments. Importantly, while this effect reduces the precision of breakpoint mapping, it does not affect the ability of the pipeline to determine the actual sequences of DIP-associated RNAs.

The third factor is the potential for base calling errors or mutations to result in erroneous junction mapping (**Fig S3B**). Even though reported junctions in this category are derived from real junctions, they can be viewed as false positives in that they are reported as distinct junctions that do not actually exist in the viral population. Altogether, these three factors set a ceiling on the maximum number of deletion junctions that can be accurately detected and mapped. Using our simulated datasets, we knew how many deletion junctions actually existed, exactly where they were located, and whether or not they were adjacent to direct repeats (see Materials and Method) that could result in incorrect mapping. This allowed us to systematically optimize the sensitivity and precision of the software pipeline.

We tested how varying the ViReMa operating parameters affected both junction-spanning read detection and actual junction reporting. We used the Cal07-200 dataset to challenge ViReMa across a range of --N parameter (number of mismatches allowed) and --X parameter (mismatch distance from the putative junction location) values. We first asked how varying the --N and --X parameters influenced the total number of junction-spanning reads detected (**Fig 2B**). We found that using N=0 (--X is irrelevant at this condition) significantly decreased the number of junction-spanning reads detected compared with non-zero --N and --X values. We next asked how increasing the --N and --X values affected the number of accurately and inaccurately mapped junctions reported (**Fig 2C**). We observed a clear correlation between the --X parameter and junction-mapping precision, as increasing the --X value decreased the number of inaccurately mapped junctions. Overall, we found that using N=1 and X=8 reduced inaccurate junction mapping to the minimum amount possible, given the occurrence of direct repeats adjacent to 23.5% (47 of 200) of junctions in the dataset.

We next asked whether setting a minimum read support cutoff (RSC) to report a junction affected the numbers of both accurate and inaccurate junctions that the pipeline identified. Requiring that a given junction be represented within a minimum number of reads can decrease the number of erroneously mapped junctions arising from base calling errors but could also result in some true junctions being lost due to insufficient read coverage. We aligned our simulated Cal07-200 dataset with Bowtie 2 and used the resulting unaligned reads to challenge ViReMa using different RSC values (**Fig 2D**). We found that the number of true junctions reported by the pipeline was very close to the theoretical maximum, with minimal drop-off across the range of RSCs tested. In contrast, we observed that the number of inaccurately reported junctions was highly sensitive to the RSC value used. An RSC of >30 was needed to lower the number of inaccurately reported junctions to the minimal limit (determined by the number of ‘fuzzy’ junctions with adjacent direct repeats in the dataset).

Altogether, these data highlight the importance of optimizing RSC values and the ViReMa --N and --X parameters for maximizing the sensitivity of junction detection while minimizing the number of false positives. We set our default values at RSC>30, --N=1, and --X=8 for subsequent analysis.

### Validation of sequencing pipeline

After optimizing the bioinformatics component of our pipeline using simulated datasets, we examined the ability of the pipeline to detect DIP-associated deletions within complex viral populations from experimental samples. Our overall strategy was based on the universal, eight segment RT-PCR approach pioneered by Zhou et al. (18). Critically, there are a number of steps within the library preparation and sequencing steps that have the potential to introduce artifacts that can compromise junction detection and analysis. In particular, we were concerned about the potential for recombination during reverse transcription, PCR, and/or sequencing to generate junctions that will be called by the pipeline (19, 20). To address this, we prepared several control sample libraries, sequenced them on the MiSeq, and ran the results through our optimized pipeline.

To quantify false positive generation during the PCR and/or sequencing steps, we constructed libraries without using actual viral RNA or reverse transcriptase. To do this, we generated an equimolar ratio mixture of full length PCR amplicons from each of the eight IAV genome segments, using reverse genetics plasmids encoding the gene segments from A/Puerto Rico/8/1934 (PR8) as templates. These amplicons were gel purified to ensure correct, full-length size, and then used as template for the universal amplification PCR and subsequent library preparation. Our analysis pipeline detected no breakpoints in this control, indicating that none of the steps in our pipeline from PCR onwards were significant sources of false positive signals.

We next sequenced a recombinant Cal07 stock that was grown under low MOI conditions to minimize the frequency of DIPs (21). We performed two independent RNA extractions and reverse transcription reactions on this stock to serve as technical replicates (named Par1 and Par2). ViReMa detected 6 and 7 DIP-associated deletion junctions from Par1 and Par2, respectively, with junction-spanning reads representing ~0.1-0.2% of the total reads **(Fig 3A)**. The majority of these reads were derived from a single shared deletion junction in HA (indicated by the following nomenclature: 615_1132_HA). 4 other DIP junctions were shared between replicates, each with low NGS read depth (ranging between 19 and 94). Two unshared junctions in Par1 and one in Par2 were actually reported in both replicates but failed to reach the level of detection in one replicate.

The significant overlap in the specific junctions that were reported from the two replicates suggested that these junctions were produced by the viral polymerase (and were thus bona fide DIP-associated sequences) rather than by the reverse transcriptase. However, the generation of the same junction in independent RT reactions could also indicate the existence of strong hotspots for RT recombination. To more directly address the potential contribution of RT-derived recombinants, we performed two independent experiments. First, we compared the junctions detected in HA segment libraries generated from Par 1 using two different RT enzymes, Invitrogen Superscript III and Agilent AccuScript. Second, we performed *in vitro* transcription of a plasmid-derived Cal07 HA segment using T7 RNA polymerase, which then was used as a template for RT-PCR to produce the amplicon library for sequencing. The IAV polymerase was not involved in this control; thus any deletions detected will have been generated by T7 polymerase or the RT enzyme.

The junctions reported from libraries generated by the two RT enzymes had significant overlap and were both dominated by 615_1132_HA (**Fig 3B**). In contrast, we detected none of the Par1-derived junctions in the library generated from T7-transcribed HA (**Fig 3B**). Although the read depth coverage was comparable to Par1, 615_1132_HA was completely absent, and the three junctions that were detected had minimal read support and were not seen in virus-derived libraries. Altogether, these results suggest that the formation of deletion junctions during the reverse transcription reaction is rare, and that the Par1-derived junctions we observed are most likely derived from true DIPs present within our viral stock, despite the stock having been prepared at low MOI. This highlights the difficulty in producing a completely DIP-free virus preparation.

**Fig 3.**
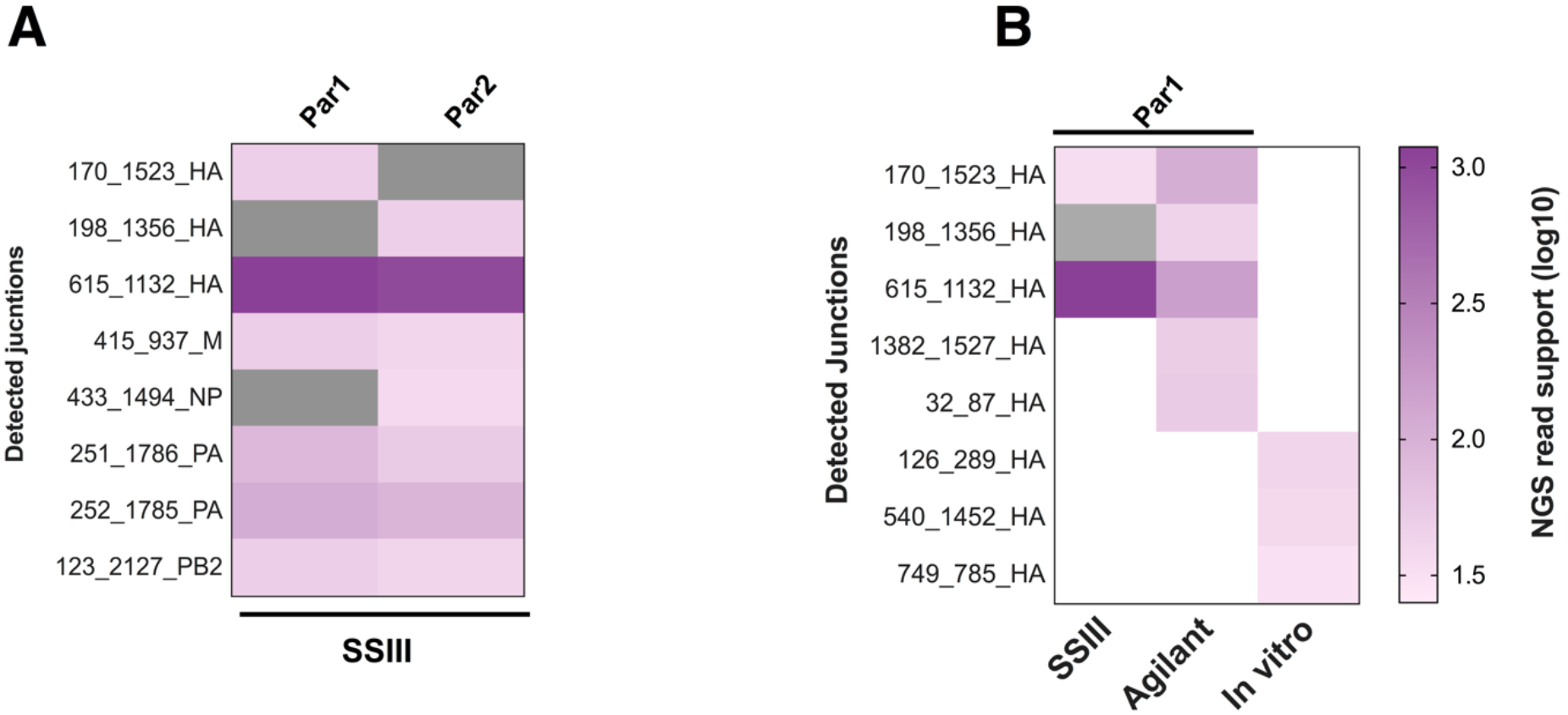
DIP-associated deletion junctions present in virus working stock. We performed two independent RNA extractions, RT reactions, PCR amplifications, and library preparations from a single recombinant Cal07 working stock grown at low MOI (Pari and Par2). **(A)** Comparison of deletion junctions detected in Pari and Par2 samples. Purple blocks represent instances where junction was detected but read support was below RSC **(B)** Comparison of HA segment junctions detected in two libraries generated from independent RT reactions using two different RT enzymes, along with a library generated from in vitro T7-transcribed viral RNA. White blocks denote no detection

### Generation of DIP-enriched populations through high MOI passage

To test the ability of the pipeline to detect real DIP-associated RNAs, we enriched for DIPs through serial undiluted passage of Cal07 in MDCK cells. We confirmed the presence of DIPs by amplifying full-length genomic cDNA at each passage and examining the size distribution of PCR products by gel electrophoresis **(Fig 4A)**, as previously described (21). The gradual disappearance of the polymerase segments, which form the majority of DIPs, and the appearance of a smear below the shortest IAV segment (NS ~0.9kb) were consistent with the accumulation of DIPs over passage. Based on these results, we picked P1, P3, and P6 as representative samples for sequencing.

We further confirmed the presence of DIPs by plotting the read coverage of the aligned reads from passages 1, 3, and 6 (**Fig 4B**). These coverage plots clearly reveal the characteristic pattern of DIP-rich populations, with much lower depth of read alignment in the middle portion of the segment compared with the termini. As expected, the number of DIP-associated deletion junctions detected by the pipeline also increased across passages, reaching the highest level at passage 6 (**Fig 4C**). To confirm that these junction-containing sequences were derived from virion rather than cellular RNA, we measured the number of reads that aligned to the host (canine) genome in our samples. We found very few reads derived from the canine genome in all the passages, compared with about 40% of the reads from RNA extracted from infected cells (**Fig S4**).

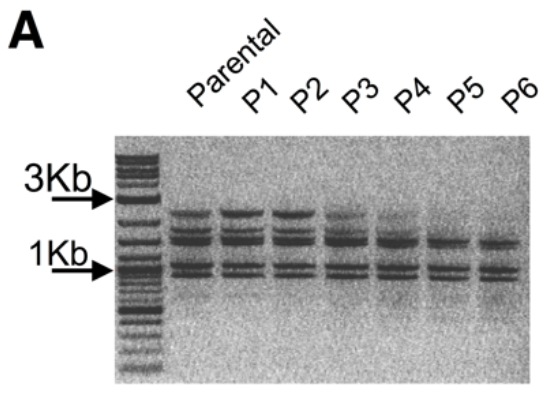

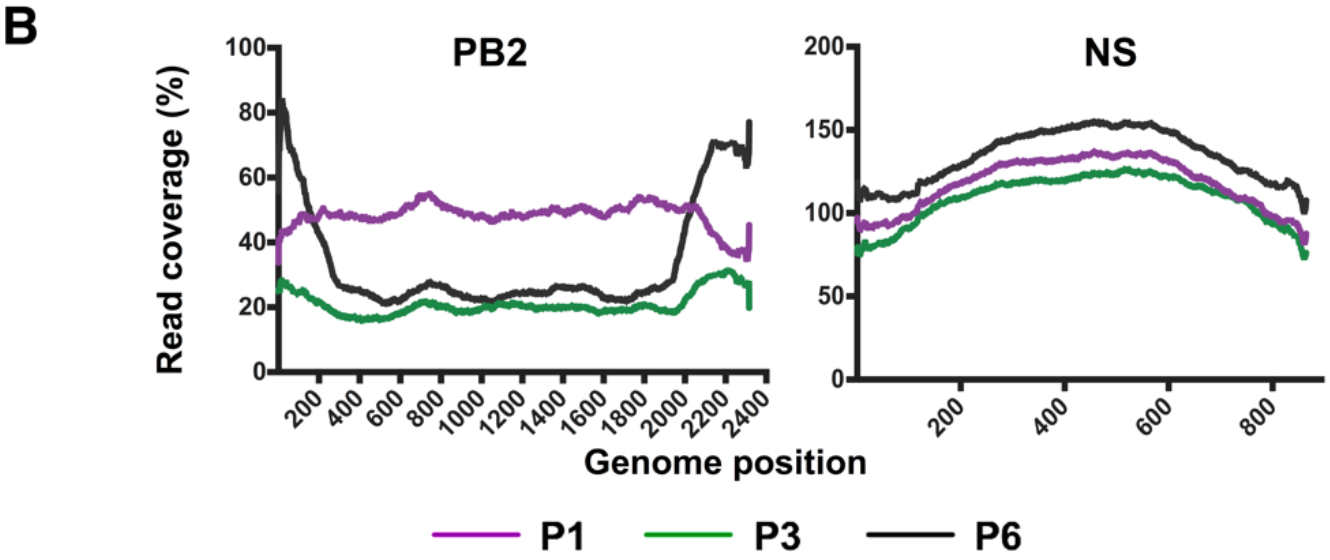

**Fig 4.**
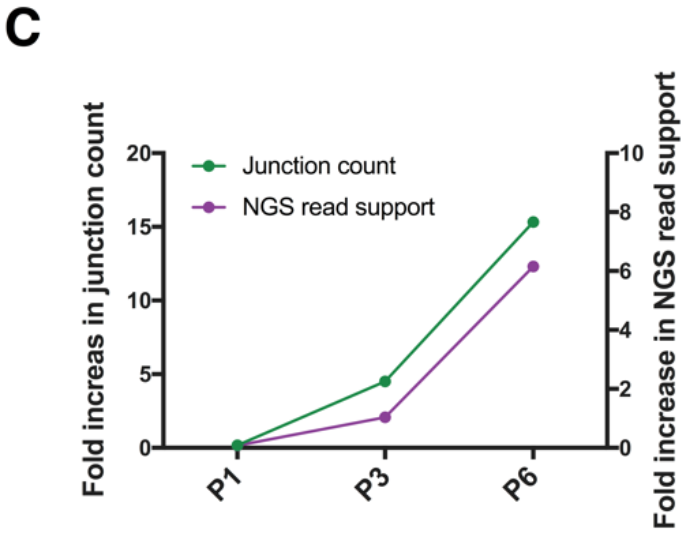
Generation of DIP-rich populations through high MOI passage. Call07 was serially passaged 6 times (P1-P6) in MDCK cells at substained high MOI. **(A)** PCR products from the indicated Cal07 populations following 8-segment whole genome amplification, visualized on a 1% agarose gel. The accumulation of delection junctions is reflected by the disappearance of the polymerase segments (~2.3kb) and the appearance of a smear below the NS segment (~0.9kb) ranging from ~0.3-0.8kb. **(B)** Coverage depth of aligned reads from the indicated passages for PB2 ans NS genome segments. The coverage was normalized to the read coverage of the parental sample (Par1). **(C)** Fold increase in the number of delection junctions (left y axis) and total read support for those junctions (right y axis) over the parental sample (par1).

### Reproducibility of pipeline performance

Multiple steps in the combined experimental/computational pipeline could introduce stochasticity into the pipeline performance, thus diminishing overall consistency and reproducibility of output. To examine the reproducibility of our pipeline’s performance, we sequenced two separate extractions of a single P6 population (Hereafter known as L1-P6-Rep1 and L1-P6-Rep2, where L refers to lineage) and compared the pipeline outputs between the two replicates (**Fig S5**). We found that the normalized read support values of individual junctions were highly correlated between the two replicate samples, whether the replicates were sequenced on the same MiSeq flowcell (Spearman R = 0.92) or separate ones (Spearman R = 0.91). Thus, the combined steps from RNA extraction to sequence analysis introduce minimal noise into the pipeline output, and pipeline performance is highly reproducible between experiments.

### Optimization of minimum read support cutoffs

Our experiments using simulated datasets revealed the importance of setting minimal RSCs for maximizing the accuracy of pipeline performance, and suggested that the optimal RSC may differ between datasets. We next attempted to optimize RSC values for our experimental dataset where we did not actually know the precise location and number of junctions present in the population (as we did with our simulated datasets). To quantify precision in junction detection for our experimental dataset, we assumed that base calling errors and mutations that result in inaccurate junction reporting would be stochastic and thus read support for these inaccurate junctions would be highly variable between technical replicates. In contrast, read support for real junctions should be consistent between replicates.

We assessed the effects of varying the RSC on the degree of correlation between junctions identified in L1-P6-Rep1 and L1-P6-Rep2. We varied the RSC values from 1 to 50 for each individual genome segment, and examined the effect on the number of reported junctions (**Fig 5**). We observed a similar pattern to that observed for our simulated data, where raising the RSC to 10 or higher resulted in a large drop-off in the number of reported junctions. We next determined the RSC value that yielded the highest degree of correlation between the two replicates. We identified distinct optimal RSC cutoff values for each segment: 20, 20, 30, 30, and 15 for PB2, PB1, PA, HA, and NA, respectively. The average of these values was used as an RSC for the remaining segments where no enough junctions were detected to perform the correlation test (see below).

We do not expect these values to be universal, as they likely are influenced by a number of factors that will vary between individual sequencing runs. Also, for different applications, it may be beneficial to lower the RSC to improve detection sensitivity at the cost of precision. Thus, we suggest running two technical replicates with each NGS run to be used as reference to establish optimal per-segment RSC values for that run.

**Fig 5.**
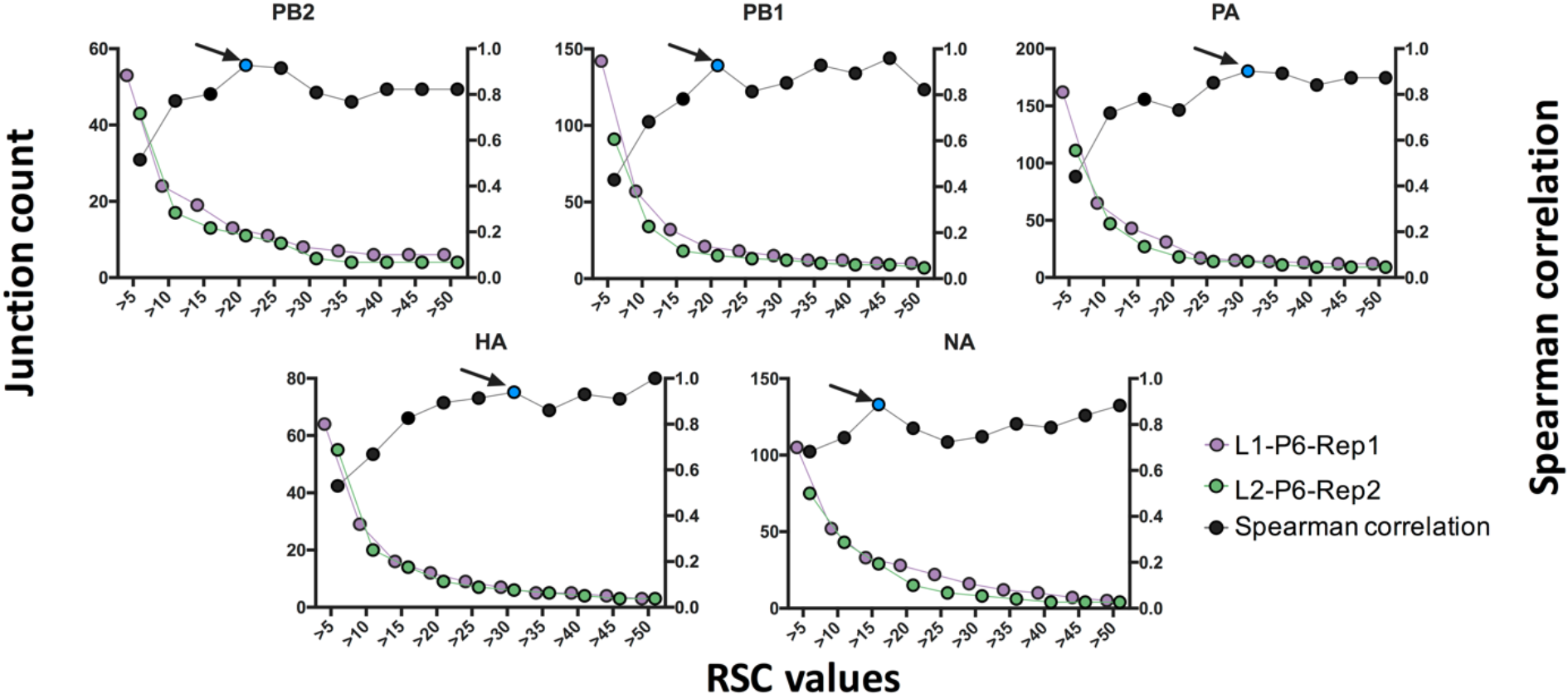
Determination of optimal read support cutoffs for experimental data. Plots showing the numbers of deletion junctions reported in the indicated genome segments for two technical library preparation and sequencing replicates generated from a single DIP-rich viral population (L1-P6-Rep1 and L2-P6-Rep2; left y axis). Black dots represent the results of Spearman correlation tests between the replicates at each RSC condition (right y axis). Blue dots indicate the point with the highest degree of correlation and minimum decrease of junction count for each genome segment.

### Analysis of DIP-associated deletion profiles

We next examined the overall diversity of DIP-associated deletion junctions within the P6 populations from the two independent lineages (L1-P6-Rep1 and L2-P6), and found dozens of distinct deletion junctions scattered across the viral genome in both lineages (**Fig 6A**). Junctions were not evenly distributed across the genome segments, as few to no junctions were detected in the NP, M, or NS segments. Within each segment, the read support for individual junctions varied significantly (**Fig 6B**). When we compared the deletion junction repertoires between the two passage lineages, we observed that a significant fraction of the detected junctions was shared between the two, and that these shared junctions exhibited a high degree of correlation in terms of read support (**Fig 6C**). These data suggest that specific DIP-associated deletions may be consistently formed by Cal07. While there was substantial diversity in terms of the number of distinct deletion junctions present, when we plotted the locations of those these junctions within the genome segments, we observed that they were largely confined within clear hotspots towards the termini of the segments with few exceptions (**Fig 6D**).

**Fig 6.**
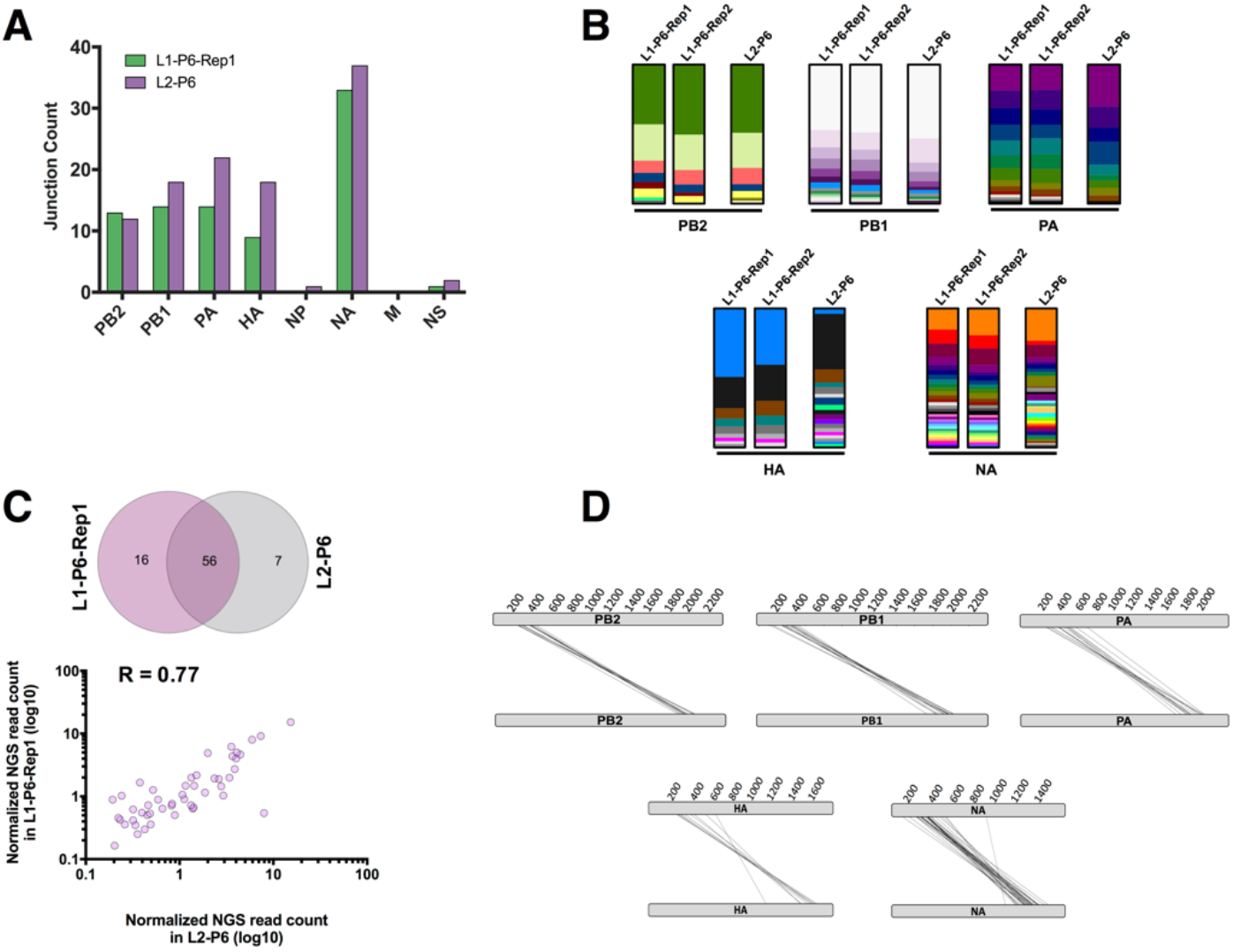
DIP-associated junctions analysis. **(A)** The total number of detected junctions in the individual genome segments for the two independent passage 6 populations. **(B)** Stacked column charts showing the proportional abundance of each deletion junction per segment between lineage 1 (including both technical replicates) and lineage 2 at passage 6. Each color bar represents a unique junction within each segment, whose height reflects the relative NGS read support, normalized to the total number of NGS junction-spanning reads for the indicated segment. **(C)** Comparison between L1-P6-REP1 and L2-P6 in relation to the number of shared junctions (upper panel) and the correlation between their NGS reads (lower panel) **(D)** Parallel coordinates diagrams showing the deletion junctions in P6-Rep1 mapped to their actual respective locations on each segment. Each individual junction is represented by a black line that connects the donor and acceptor sites of the breakpoint.

### Effect of varying template input on pipeline performance

We next asked whether the amount of cDNA template that goes into the library preparation affects the sensitivity and stochasticity of junction detection by the pipeline. We serially diluted both the amount of viral RNA template used in the RT reaction and the amount of cDNA template used in the PCR and compared pipeline outputs from the DIP-rich L1-P6-Rep1 population. We first tested the correlation of detected DIP-associated junctions between a limited number of dilutions ranging from 1:3 to 1:15. We observed that the correlation of read support values between specific junctions across dilutions was more consistent when cDNA was diluted, rather than RNA, suggesting that RNA dilution may increase the stochasticity of downstream PCR amplification (**Fig S6A**).

Based on this, we performed whole genome PCR using a dilution series of L1-P6-Rep1-derived cDNA (spanning roughly 4*10^8^ to 4*10^6^ NP genome equivalents per PCR) as template (**Fig 7A**). We observed that there is an optimal amount of input cDNA template for maximizing junction detection. Diluting the input cDNA 1:120 (corresponding to ~4*10^6^ NP genome equivalents) increased the number of detected junctions over 4-fold compared with undiluted input. Although the number of DIP-associated junctions was increased, the distribution of junctions across segments and their mapped locations were consistent with our earlier results (**Fig S6B and Fig 6**).

Further dilution of input template beyond 1:120 resulted in a decrease in sensitivity. Importantly, dilution across the range tested did not result in a failure to detect any of the junctions reported in the undiluted sample. We also observed that the correlation of read support values between specific junctions across dilutions tracked closely with the sensitivity (**Fig 7B**). Altogether, these observations indicate that optimization of the cDNA template input amount can significantly improve the sensitivity of DIP-associated junction detection.

**Fig 7.**
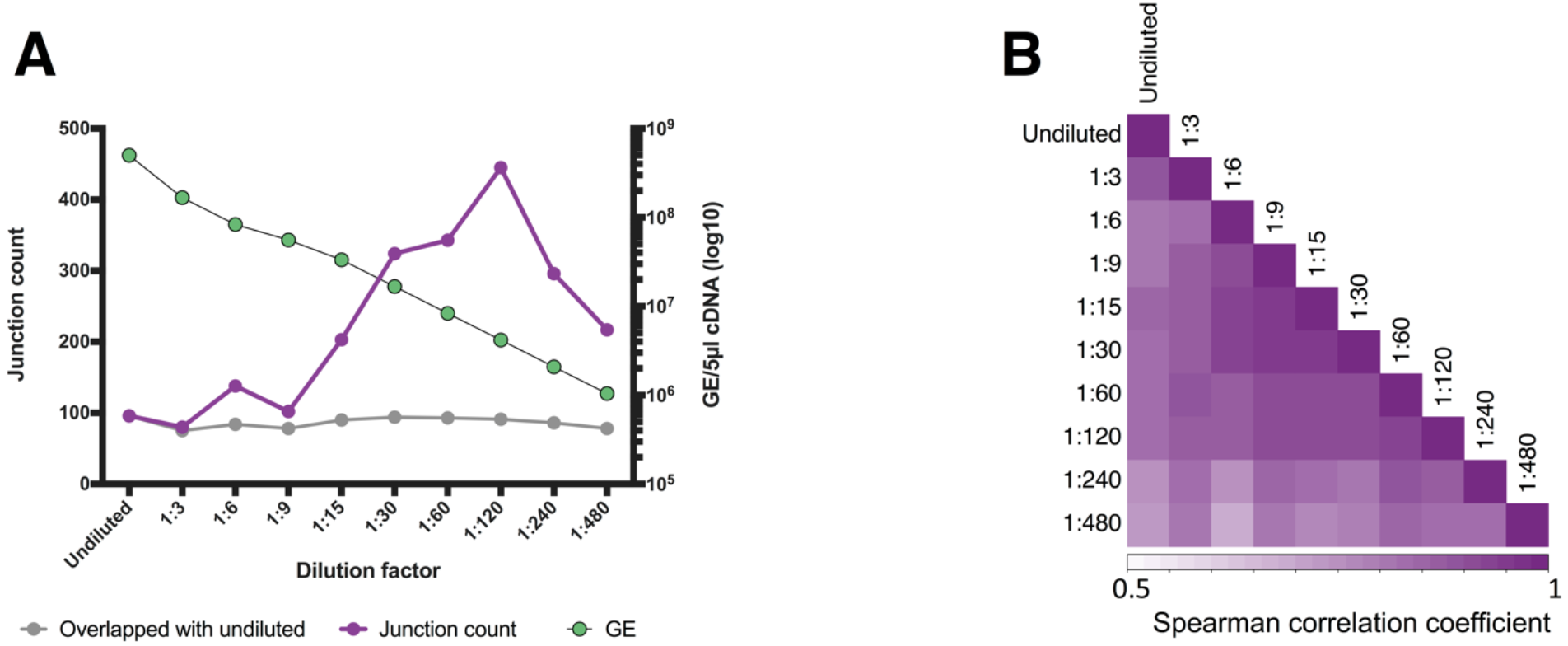
Effects of viral template input on the detection of DIP-associated junctions. We serially diluted cDNA generated from the L1-P6-Rep1 sample, and compared sequencing results between libraries generated with these dilutions as templates. **(A)** For each dilution, the total numbers of detected junctions (purple) are shown, along with the number of specific junctions detected that were also detected in the undiluted sample (grey). The copy number of viral cDNA molecules included in downstream PCR and library preparation for each dilution was determined by RT-qPCR (green; right y axis). **(B)** Read support values for all deletion junctions common across the diluted and undiluted samples were normalized to the total number of deletion junction-spanning reads for each sample and used to perform a Spearman correlation between all pairs of samples using R cor function.

### Lack of association between direct repeats and junction formation

Direct repeat sequences (detailed in **Fig S2 and S3A**) are common across the IAV genome and have previously been hypothesized to contribute to DIP-associated deletion formation by promoting viral polymerase slippage (10, 15). We leveraged the large number of DIP-associated deletion junctions that we identified in this study to test this hypothesis. We asked whether the deletion junctions in the DIP-enriched sample L1-P6-Rep1 were found more frequently adjacent to direct repeats than would be expected if the junctions were located randomly in the viral genome. We compared the frequency of deletion junctions associated with direct repeats between the L1-P6-Rep1 and L2-P6 populations (where all deletions are formed by the viral RdRp) and the Cal07-200 simulated dataset, where all deletions are randomly localized (**Fig 8**). The frequency of direct repeats of varying lengths at junction sites in the real viral populations was not significantly different than that seen in the simulated data, indicating that direct repeat sequences are not enriched at DIP-associated junctions and arguing against a significant role for direct repeats in DIP formation.

**Fig 8.**
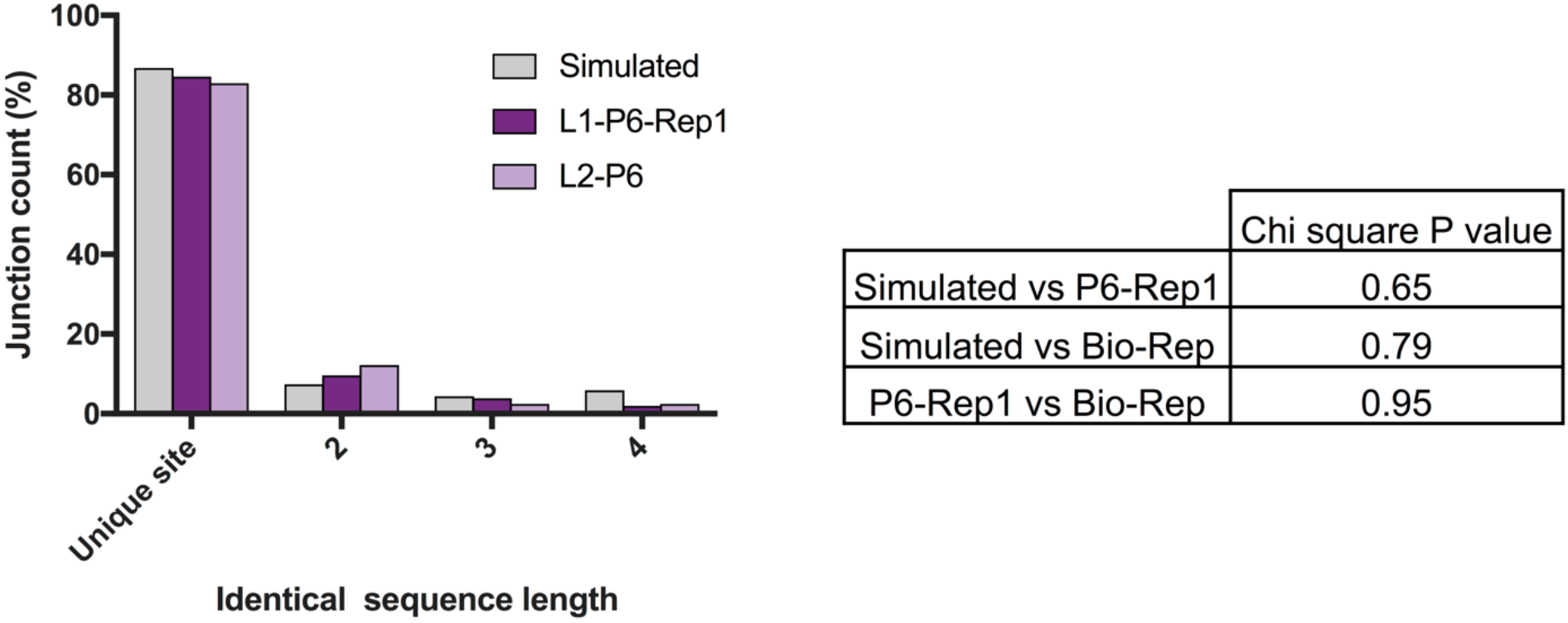
Direct repeat sequences are not over-represented at DIP-associated deletion junctions. The percentages of deletion junctions within the polymerase segments that occurred at unique sites or at sites with direct nucleotide repeats with lengths 2-4nts was compared between L1-P6-Rep1, L2-P6, and the Cal07-200 simulated dataset. The number of junctions was plotted and compared by Chi Square. The table shows the Chi Square P values between every possible pair of the samples.

## DISCUSSION

Sensitive and accurate detection of DIP-associated sequences within viral populations is critical for defining how DIPs form and function during IAV infection. Here we outline a pipeline to detect DIP-associated junctions within viral populations using Illumina-based short read sequencing, and validate its performance using a combination of simulated and experimental control datasets.

Our primary goal was to develop and optimize a reasonably simple and straightforward sequencing framework that accounts for the potential artifacts that can potentially confound NGS-based DIP detection efforts. We chose the Illumina sequencing platform because it is widely available, easy to use, conducive to sample multiplexing, and because it has a relatively low rate of base-calling errors. One concern that we had initially was that recombination during reverse transcription, PCR, or sequencing might make identification of bona fide DIP-associated junctions a challenge. Two recently developed technologies, CirSeq and ClickSeq, largely eliminate this issue, but also significantly increase the amount of labor involved in library preparation (22, 23). We observed that the occurrence of non-viral recombination that occurs during our library preparation and sequencing procedures was vanishingly small, and can effectively be ignored. Thus, while both Cirseq and ClickSeq are enormously useful in certain circumstances, our data indicates that such methods are not required to generate highly accurate and sensitive profiles of IAV DIPs.

A significant shortcoming of the method we detail here is that the measured read support for individual deletion junctions does not necessarily reflect the actual frequencies of these deletions within the viral population. This is due to both biasing of PCR amplification towards shorter products, as well as the uneven distribution of read coverage across the viral genome. For situations where the accurate measurement of individual DIP genotype frequencies is critical, we recommend pairing a cDNA barcoding method such as primer ID (24, 25) with a platform capable of long-read sequencing, such as PacBio or Oxford Nanopore (26). Alternatively, direct sequencing of viral RNA using the Oxford Nanopore platform may also prove to be useful for accurate measurement of junction frequencies (27).

When we used our pipeline to examine DIP-enriched viral populations generated through serial high-MOI passage, we detected dozens of distinct DIP-associated deletion junctions, revealing a high degree of diversity within the DIP population. Although the majority of these DIP-associated junctions were derived from the polymerase segments as expected, we also detected a substantial proportion of deletions within the HA and NA segments, but not the NP, M, and NS segments. The non-random distribution of junctions across the genome segments mirrors what has been reported elsewhere, and highlights how little we know about the specific molecular mechanisms that regulate DIP formation.

We hope that the approach detailed here, and the associated bioinformatics pipeline prove useful to other groups interested in defective interfering particle biology. Our approach is optimized for influenza virus sequences; however, the approaches and controls detailed here can easily be adapted to other RNA virus systems.

## MATERIALS AND METHODS

### Viruses and Cells

Madin-Darby canine kidney cells (MDCK; obtained from Dr. Jonathan Yewdell) and human embroyonic kidney 293 cells (293T; obtained from Dr. Joanna Shisler) were grown in Minimal Essential Medium (MEM) + GlutaMAX (Gibco), supplemented with 8.3% fetal bovine serum (Seradigm), at 37°C and 5% CO2. Recombinant A/California/07/09 (Cal07) virus was rescued via the standard 8-plasmid reverse genetics approach. Briefly, 60-90% confluent 293T cells were transfected with 500ng of the following plasmids (pDZ::PB2, pDZ::PB1, pDZ::PA, pDZ::HA, pDZ::NP, pDZ::NA, pDZ::M, pDZ::NS) using JetPRIME (Polyplus) according to the manufacturer’s instructions. Cal07 reverse genetics plasmids were originally obtained as A/California/04/2009-encoding plasmids from Dr. Jonathan Yewdell. We introduced A660G and A335G substitutions into the HA and NP plasmids, respectively, to convert them to match the amino acid sequence of A/California/07/2009 HA and NP (NCBI accession# CY121680, CY121683). A seed stock was prepared by amplifying a plaque isolate from the rescue supernatants. Virus working stocks were generated by infecting MDCK cells with seed stock at an MOI of 0.001 TCID50/cell and collecting and clarifying supernatants at 48 hpi.

### Generation of DIP stocks through high MOI passage

Confluent MDCK cells in 96-well plates were infected with IAV Cal07 at an MOI of 5 TCID50/cell. Supernatants (200*μ*l total per well) were harvested at 24 hpi (passage 1) and pooled. 100*μ*l/well of this pooled supernatant was used to infect a 96 well plate of fresh MDCK cells to generate the next passage. This process was repeated 6 times to produce passages 1-6 in two independent lineages (1 96-well plate per lineage).

### IAV Genome amplification

Viral RNA was extracted from 140*μ*l of cell culture supernatant using the QIAamp viral RNA kit (Qiagen) and eluted in 60μl distilled H2O (dH2O). For cDNA reactions, 3*μ*L of RNA was mixed with 1*μ*L (2*μ*M) MBTUni-12 primer (5’-ACGCGTGATCAGCRAAAGCAGG-3’) + 1*μ*L (10*μ*M) dNTPs + 8*μ*L dH2O. The mixture was incubated for 5 minutes at 65°C and then placed on ice for 2 min. Subsequently, the mixture was removed from ice and the following was added: 1*μ*L SuperScript III RT (Invitrogen), 4*μ*L of 5X First-Strand Buffer (Comes with SSIII kit), 1*μ*L of DTT, 1*μ*L RNase-in (Invitrogen). The reaction was incubated at 45°C for 50 min, followed by a 15 min incubation at 70°C for inactivation. 5*μ*L of cDNA product was mixed with the following for PCR amplification: 2.5*μ*L (10*μ*M) MBTUni-12_4R primer (5’-ACGCGTGATCAGCRAAAGCAGG-3’), 2.5*μ*L (10*μ*M) MBTUni-13 primer (5’-ACGCGTGATCAGTAGAAACAAGG-3’), 0.5*μ*L Phusion polymerase (NEB), 10*μ*L - 5x HF buffer, 1*μ*L (10mM dNTPs mix), and 28.5*μ*L dH2O. The PCR reaction conditions used: 98°C (30 s) followed by 25 cycles of 98°C (10 s), 57°C (30 s) and 72°C (1:30 min), a terminal extension of 72°C (5 min), and a final 10°C hold. PCR products were purified using the PureLink PCR purification kit (Invitrogen) with the <300nt cutoff option and eluted in 30*μ*L dH2O. There was no difference in deletion junction detection when we purified the PCR products with the lower cutoff option (data not shown).

### NGS library preparation

We started with ~20ng of the PCR products in a volume of 50*μ*l. The Covaris M220 sonicator (Covaris) was used to fragment the DNA. Three different conditions were used to generate different average fragment lengths of 300, 500, 700 base pairs (bp): (I) 300 bp = Peak Power 50, Duty Factor 20 and Cycles/Burst 200 for 2:40 min, (II) 500 bp = Peak Power 50, Duty Factor 10 and Cycles/Burst 200 for 1:30 min, and (III) ~600 bp = fragment length Peak Power 50, Duty Factor 10 and Cycles/Burst 200 for 1 min. In our hands, the fragmentation length did not have any effect on our sequencing results (data not shown). For the sake of consistency, we used the 300 bp fragmentation length. To confirm the PCR products, we visualized the amplicons on a Fragment Analyzer (AATI) with the DNF-486 high sensitivity NGS kit before and after fragmentation. Next, we used KAPA Hyper Prep kit (Roche) to construct the libraries according to the manual. To eliminate the possibility of index hopping (or index switching), we used the TruSeq Unique Dual Indexes (UDI) from Illumina. The Adapter ligation step was carried out with 5*μ*l of Truseq UDIs diluted 1:10 with 10nM Tris. For maximum efficiency we increased the ligation time to 30mins. We then performed 3 cycles of PCR with the Kapa library amplification primers diluted 1:5 in water followed by a cleanup step with 40*μ*l of AxyPrep Mag PCR beads (Thermofisher). We then mixed the libraries at an equimolar ratio and carried out a qPCR to accurately quantitate the library pool and maximize the number of clusters in the sequencing flowcell. A size selection step was not needed. Finally, the pooled libraries were sequenced with paired-ends 2×250nt reads on an Illumina MiSeq using V2 chemistry. The fastq files were generated and demultiplexed with the bcl2fastq v2.20 Conversion Software (Illumina).

### Simulated Datasets

All the simulated datasets used in this study were generated by MetaSim (v0.9.1) (28), a genomic and metagenomics simulator. Several reference library sequences composed of WT reference sequences of IAV Cal07 or PR8 (see Table S1 for NCBI accession numbers), mixed with a defined DIP sequence population - generated randomly within the first and last 600 nts of all the segments - were used in Metasim for data simulation. The configurations were fixed across all datasets to maintain the preferable conditions. The reference sequences were fragmented into 350 nts fragments length with a standard deviation of +/− 50 and were simulated into ~1 million 2×250nts paired-end reads per sample, with a total mutation rate of ~1 % based on the published Illumina empirical error model, and corresponds to substitutions as the indel error rate is negligible within Illumina MiSeq. One dataset was simulated with no DIP sequences as a control sample for any computational artifacts. Metasim generated two FASTA files of 1 million reads per file per sample (~2 million single-end reads = 1 million paired-end reads), which subsequently were used for the optimization process.

### Sequencing analysis of DIP-associated junctions

The raw sequencing reads were quality-filtered by Trimmomatic (v0.36) (Parameters: `ILLUMINACLIP:TruSeq3-PE-2.fa:2:15:10 SLIDINGWINDOW:3:20 leading:28 trailing:28`) (29) and any reads shorter than 75nts were removed from the datasets. The paired reads were concatenated into one file and treated as single-end when aligned end-to-end to the WT reference sequences using Bowtie2 (v2.3.1) (Parameters: --score-min `L,0,−0.3`). Subsequently, the algorithm ViReMa (v0.10) was used to analyze the remaining un-aligned reads (putative junction-spanning reads) (Parameters `-DeDup --MicroInDel_Length 20 --Defuzz 3 --N 1 --X 8`). Next, the DIP-associated deletion junctions and their read support were extracted from ViReMa output files and sorted per segment, using an in-house Perl scrip, for data analysis and visualization. To detect any MDCK genome leakage, the datasets were aligned against the dog genome (assembly CanFam3.1). All scripts are available at *https://aithub.com/BROOKELAB/Influenza-virus-DI-identification-pipeline*.

### Quantification of sensitivity and precision

To calculate the actual number of junction-spanning reads in Fig 2A, reads that derived from DIP-associated sequences were counted by their FASTA headers, which contain the source of each read, produced by MetaSim. To calculate the maximum theoretical sensitivity of ViReMa (Fig 2B) based on seed length of 25nts and two allowed mutations (--N = 2), the number of mutations was subtracted from the seed length, which on its turn was multiplied by 2 to account for both termini ((25-2)*2=46). Subsequently, this number was subtracted from the possible cutting site of a 250nts read and divided by the total number of cutting sites and multiplied by 100 ((249-46)/249*100)=81.5%). To calculate the number of accurately and inaccurately mapped junctions in Fig 2C,D, the seed sequences of the Cal07-200 dataset were used against ViReMa with --N set to 0, and the remaining parameters were kept the same. These sequences were generated initially to establish the seed for MetaSim to simulate sequencing, therefore their lengths are varied between ~350-1800. The long ones were trimmed to <1000nts, so ViReMa would take them as reads (the maximum default read length that ViReMa could take is ~1024nts) and, critically, the junction locations were maintained. The junctions that occurred within the first or last 25nts were removed (4 junction sequences). Finally, the junctions that accurately mapped were counted, which found to be 149 versus 47 inaccurately mapped junctions.

### Correlation analysis

For the correlation tests, the NGS read support count for each DIP-associated junction was normalized to the total detected junction-spanning reads of every sample. Next, the correlation was calculated based on Spearman rank correlation using either R (cor function) or an online tool at: http://www.biostathandbook.com/spearman.html

## ACKNOWLEDGMENTS

We are grateful to other members of the lab for helpful comments and critical reading of the manuscript, as well as to Dr. Alvaro Hernandez and the staff at the high-through sequencing and genotyping unit within the Roy J. Carver Biotechnology Center for excellent advice and technical assistance. This work was generously funded by the Defense Advanced Research Projects Agency under contract DARPA-16-35-INTERCEPT-FP-018.

